# Percolation models of pathogen spillover

**DOI:** 10.1101/497750

**Authors:** AD Washburne, D Crowley, D Becker, K Manlove, R Plowright

## Abstract

A series of logical events must occur for a pathogen to spill over from animals to people. The pathogen must be present in an animal reservoir, it must be shed from the reservoir into the environment or be transferred from the reservoir to a vector, it must persist in the environment or vector until contact with a human or amplifier host, and it must successfully enter, colonize, and reproduce within the human. These events each represent a barrier the pathogen must cross to successfully infect a human. Percolation models of pathogens completing the series of barriers or logical events can connect models of spillover risk with standard tools for statistical inference.

Here, we develop percolation-based models of spillover risk and a theoretical framework for managing spillover as an inextricably multilevel process. Through analysis and simulation, we show that estimated associations between level-specific covariates and spillover events will err towards associations from dominant pathway to spillover, a potential problem if there are alternative pathways to spillover with different associations with covariates. Furthermore, estimated associations between covariates and spillover will better reflect associations between covariates and success probabilities of bottleneck events with the highest pathogen attrition rates in the data observed. If one agrees with a percolation model for spillover, then GLMs should not be used to estimate relative importance of various levels. We recommend always using nonlinear models for predicting spillover risk with quantitative covariates and discuss why switching regression models may be well suited for avoiding some obvious pitfalls in predicting spillover from alternative pathways or wildlife reservoirs. Finally, we demonstrate how percolation models formalize an intuitive management paradigm for mitigating risk in the inherently multilevel process of pathogen spillover.

## Introduction

A series of logical events must occur for a pathogen to spill over from animals to people. The pathogen must be present in an animal reservoir, it must be shed from the reservoir into the environment or be transferred from the reservoir to a vector, it must persist in the environment or vector until contact with a human or amplifier host, and it must successfully enter, colonize, and reproduce within the human. Predicting pathogen spillover by understanding the processes shaping these various logical events is a major challenge for contemporary epidemiology [1, 2, 3].

All predictions are based on mathematical models, yet one of the great challenges of mathematical biology, absent the laws and governing dynamics present in physics and chemistry, is deciding on a model of suitable complexity; in the words of George Box, “all models are wrong but some are useful” [4]. A general space of all possible spatiotemporal models for pathogen spillover has been fleshed out in a recent paper [3]. However, linking general dynamical systems of abstracted functional forms connecting everything from wildlife infection d dynamics and pathogen shedding to dose-response curves may fall short of predicting spillover risk due to the inherent complexity of abstract, deterministic models of everything.

Here, we describe a series of probabilistic models based on the percolation of pathogens through a series of logical steps between reservoir host infections to downstream human infections. Percolation models of pathogen spillover adhere to biological first principles and still allow tractable results on how to infer the relative importance of various processes in pathogen spillover. By allowing tractable results and a tunable level of complexity, our probabilistic framework may be more useful for quantifying spillover risk and the uncertainty about our predictions.

Percolation processes in mathematics are the stochastic movement of material on graphs [5]. The pathway to spillover could be modeled as a directed graph with nodes representing various pools or measurement points (reservoir hosts, environment, etc.), and edges representing the potential movement of pathogens between those pools. In such a graph, one can tally pathogen loads at the nodes and model probabilities of successful passage along edges.

When a pathogen is shed, it enters a pool of pathogens in the environment. When it survives long enough in the environment to contact a human, the shed pathogen enters a pool of pathogens available to infect a host. As the logical events on the pathway to spillover occur, pathogens from animal reservoirs move along various states or pools on the pathway towards infecting a human. Percolation models of pathogen spillover can connect the randomness of real-world data – often count processes of spillover events – with statistical models of the series of logical events.

This paper has two parts. First, we enumerate different percolation model structures corresponding to different assumptions about the pathway(s) of pathogen spillover. In the context of model structures, we develop conceptual tools useful for discussing spillover risk and assignment of relative importance to various steps of the spillover process. The model structures illuminate why log-probabilities yield a more natural way to analyze such percolation processes, yielding some important concepts for management such as the amount of variance in spillover risk that is manageable. While log-probabilities are useful, they introduce a nonlinearity which provides insight into potential pitfalls of statistical inferences of spillover risk and its associations with covariates.

Second, we examine parameter estimation in percolation models. Percolation processes provide tractable analysis of rate parameters at various pools in the pathway to spillover. However, true rates are often nonlinear in the natural or canonical parameters one infers via regression on counts of spillover events through generalized linear models (GLMs) [6]. When attempting to infer associations between covariates and the rate of pathogen spillover from multiple reservoirs or multiple pathways, the results will be biased towards those of the dominant reservoir or pathway.

Percolation-based models of pathogen spillover provide easy visualizations of model structure, a tractable route for analysis of spillover risk, agreement with first principles of pathogen spillover, conceptual tools for management, and a clear connection between model structure and statistical inference.

## Model Structures

A series of logical events must occur for a pathogen to spill over from wildlife to people. In this paper, we consider an integer number, *X*, of pathogen particles released from the reservoir and surviving a series of logical events leading to *Y* spillover events. These percolation model structures can be illustrated with a graph indicating the pathway to pathogen spillover. For example, the graph below, read from left to right,

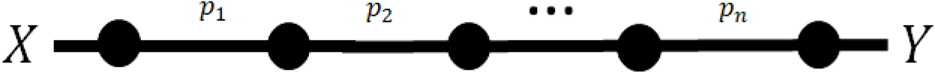

represents *X* pathogens produced from wildlife reservoirs. The pathway to pathogen spillover is partitioned into a series of *n* logical events with probabilities *p*_1_, *p*_2_, … *p_n_* of occuring. At the end of the graph, we denote a total number of spillover events, *Y*. Under this model, spillover only occurs if each of the n steps in the progression occurs. Such a graph could be used to represent a percolation model for the pathway to spillover of Nipah virus, for example [Oppenshaw et al., this issue]. We refer to this as the “serial” model for pathogen spillover.

Another scenario is a case in which, for one part of the pathogen spillover pathway, there exist alternative pathways. For example, Ebola virus can spill over directly from bats to people through bushmeat hunting [7] but spillover is more likely to occur through contact with the infected carcass of an amplifier host such as a forest antelope or non-human primate [8]. The percolation model for Ebola virus will have the graph structure

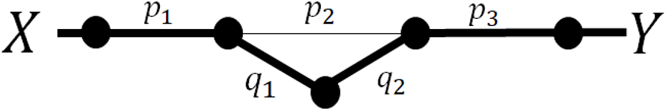

where the thickness is used to illustrate the dominant pathway and the route to the amplifier has two new probabilities: *q*_1_, the probability of infecting an amplifier, and *q*_2_, the probability an infected amplifier contacts a human with an infectious dose. The percolation models we investigate below are summarized in Table 1.

**Table 1.**
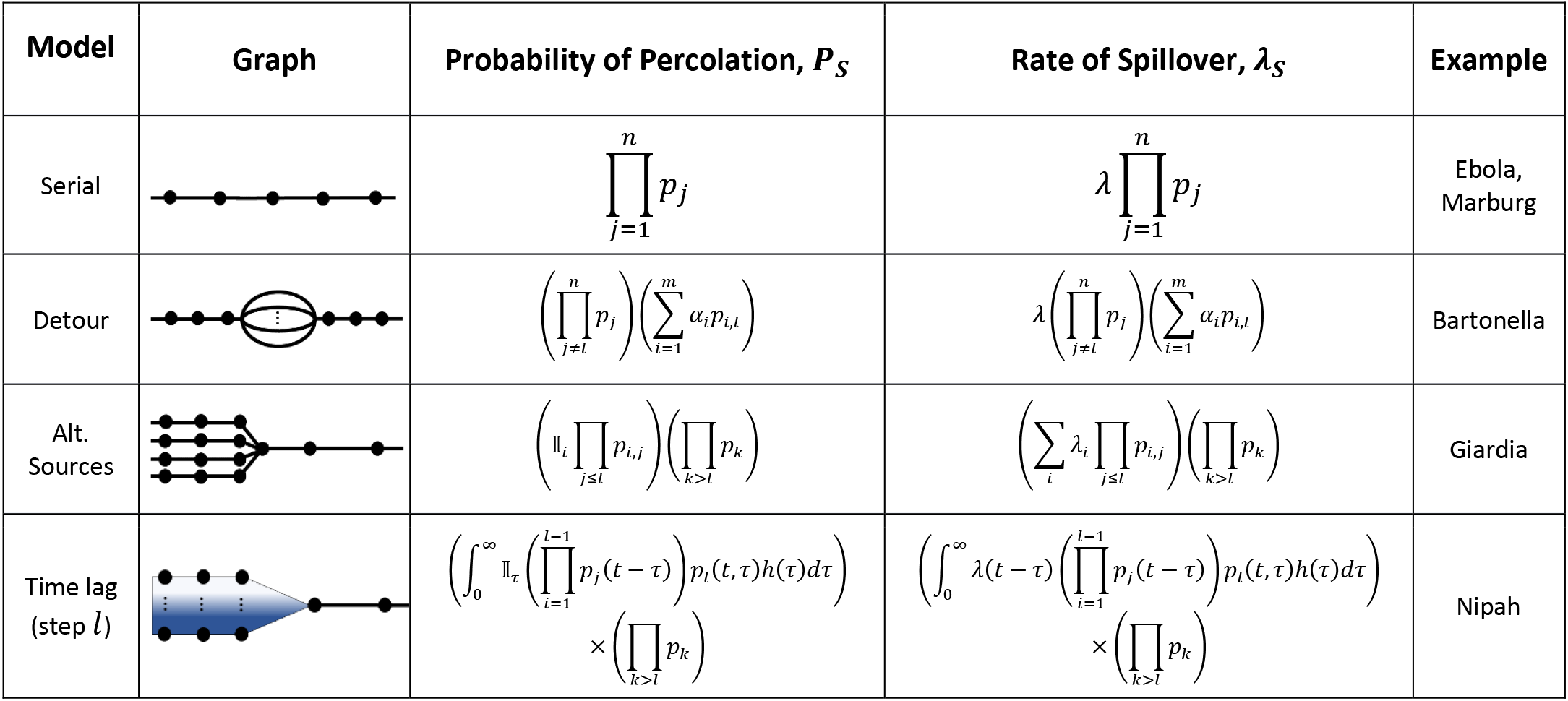
Percolation-based models of pathogen spillover allow graphical visualizations of model structure (the graphs) and a clear connection between a multilevel model and resulting probabilities of percolation and rates of spillover. Here, *λ*_*i*_ is the rate of shedding from source *i, p_j_* is the probability of surviving level *j, p*_*i,j*_ the probability of surviving level *j* along pathway *i*, and 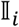 is the indicator function equal to 1 if the pathogen is shed from source *i* and 0 otherwise. Percolation models can be easily adapted for different pathogens with different pathways to spillover, and most adaptations yield similar nonlinearities whose impacts on spillover rates and statistical inference we discuss in this papers.

| Model | Graph | Probability of Percolation, $P_S$ | Rate of Spillover, $\lambda_S$ | Example |
| --- | --- | --- | --- | --- |
| Serial | | $\prod_{j=1}^n p_j$ | $\lambda \prod_{j=1}^n p_j$ | Ebola, Marburg |
| Detour | | $\left( \prod_{j \neq l} p_j \right) \left( \sum_{i=1}^m \alpha_i p_{i,l} \right)$ | $\lambda \left( \prod_{j \neq l} p_j \right) \left( \sum_{i=1}^m \alpha_i p_{i,l} \right)$ | Bartonella |
| Alt. Sources | | $\left( \mathbb{I}_i \prod_{j \leq l} p_{i,j} \right) \left( \prod_{k > l} p_k \right)$ | $\left( \sum_i \lambda_i \prod_{j \leq l} p_{i,j} \right) \left( \prod_{k > l} p_k \right)$ | Giardia |
| Time lag (step $l$) | | $\left( \int_0^\infty \mathbb{I}_\tau \left( \prod_{i=1}^{l-1} p_j(t-\tau) \right) p_l(t,\tau) h(\tau) d\tau \right) \times \left( \prod_{k > l} p_k \right)$ | $\left( \int_0^\infty \lambda(t-\tau) \left( \prod_{i=1}^{l-1} p_j(t-\tau) \right) p_l(t,\tau) h(\tau) d\tau \right) \times \left( \prod_{k > l} p_k \right)$ | Nipah |

For notation, we will always use *X* as the random variable representing the number of virions shed or released from the reservoir and *P_S_* – the probability of spillover - will denote the probability that an individual pathogen particle infects a human or other recipient host. We use *p* to denote probabilities for logical events on the pathway to pathogen spillover, with single subscripts (*p_j_*) representing the *j^th^* event and dual subscripts (*p_i,j_*) representing the *i^th^* alternative pathway for the *j^th^* event. Finally, *Y* will represent a filtered number of particles after a series of logical events, with *Y_j_* representing the number of pathogen particles surviving *j* events. In general, the subscript *j* will denote event *j* in a series of events to pathogen spillover, *i* will indicate an alternative pathway, and *l* will be reserved for the counts of pathogens surviving up to some intermediate node or measurement pool in the pathway to spillover.

### Serial (Figure 1) – conceptualization of percolation

**Figure 1.**
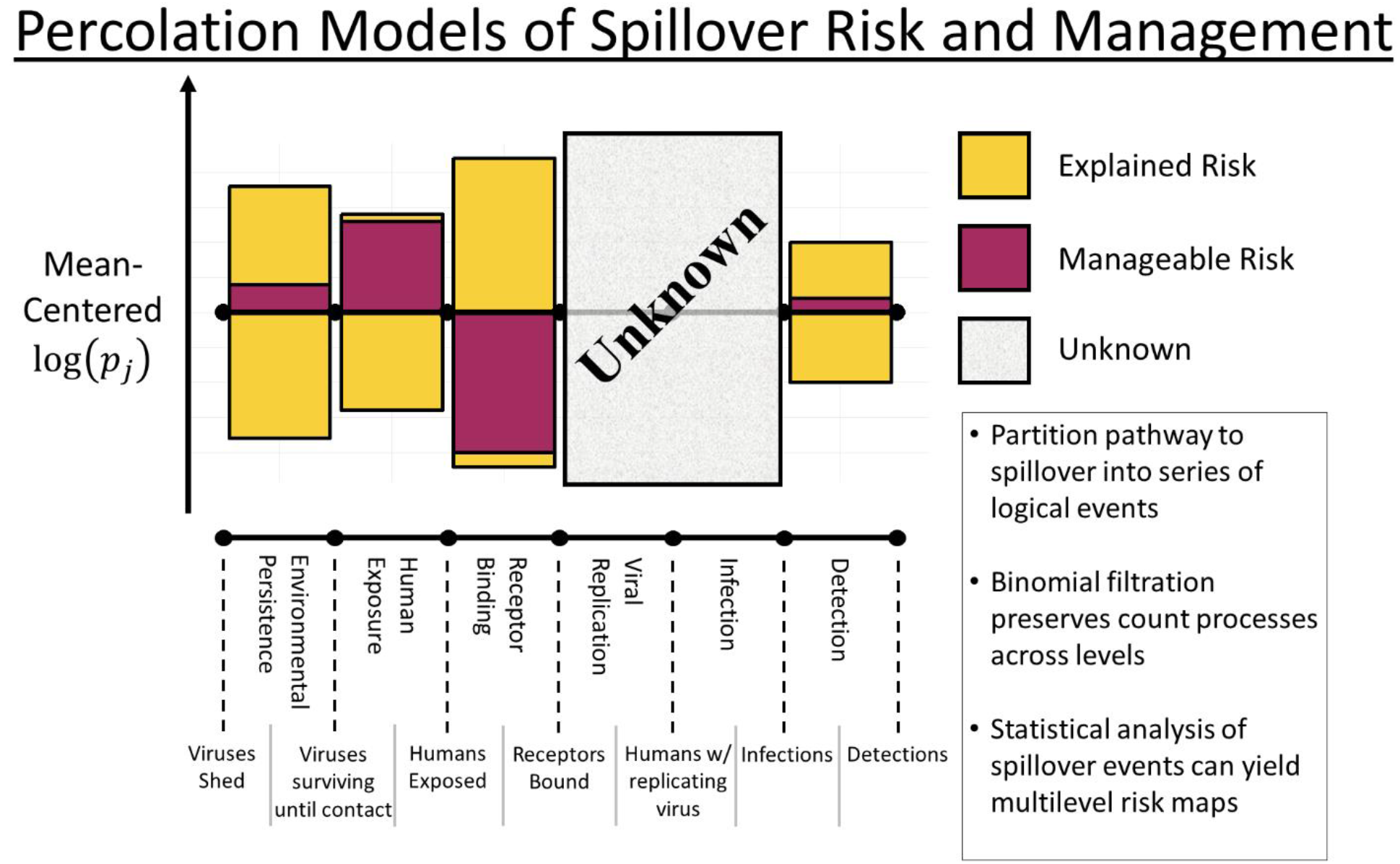
Partitioning the pathway to spillover into a series of logical events allows one to model, study, conceptualize, and visualize risk maps of spillover across levels. Percolation models subdivide the overall probability of spillover into a series of events, such as environmental persistence or viral replication, in-between which there are pools of potential observations, such as the number of human exposures or the number of infected humans. Some variation in the overall risk of pathogen spillover can be explained through known associations between log-probabilities log(*p*_*j*_) of each event *j* in the series, and external covariates. A management action can differentially impact the probabilities of each event happening, leading to a degree of manageable risk under a proposed management regime. Covariances between attrition rates at different levels, and even directionality of changes following a management action, can be visualized with asymmetric graphs of manageable risk. Pathogen spillover is an inextricably multilevel process, and as such quantifying the manageable risk requires knowing the impacts of management actions on every one of the series of events happening. Unknown or unstudied levels can impact the overall effect of an action on spillover risk, and such unknown effects must be explicitly recognized and can either assumed to be unaffected or prioritized for further study.

The simplest percolation model for pathogen spillover is to partition a single pathway into *n* logical events, *j* = 1, …, *n*, each with probability *p_j_* of occurring given all previous events have occurred. One way to formalize this model is with a series of Bernoulli random variables *B_j_*) with probability of success *p_j_*, indicating whether or not a pathogen makes it through level *j* on the pathway to spillover. The random variable *B_S_* = Π_*j*_*B_j_* indicates whether or not a pathogen makes it through every level to infect a person. In this serial model of a single pathway to pathogen spillover, the probability a given infectious particle causes a human infection is

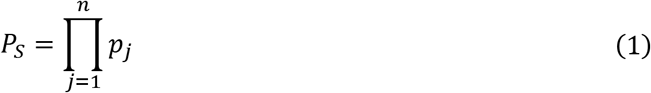

Percolation processes operate well with count processes frequently used to model infectious particle release or spillover events. If we assume the number of viruses released into the environment is a Poisson random variable, *X~Pois*(*λ*), and a serial percolation process filters viruses – each virion having probability *p*_*j*_ of surviving step *j* on the pathway to spillover – then the number of spillover events is

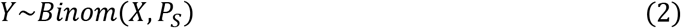

which is easily shown to follow a Poisson distribution, *Y~Pois*(*P*_*S*_*λ*). Similarly, if *X~NegBinom*(*μ,ψ*), then one can show *Υ~ΝegΒinom* (*Ρ*_*S*_*μ, ψ*) (see supplemental information for proofs). The stability of count distributions filtered through percolation models is a useful feature for analysis and statistical inference as it connects shedding rates and survival probabilities to the end result of a random number of spillover events.

Some intuition about the behavior of percolation models of pathogen spillover can be obtained by taking the logarithm of equation 1. If each *p*_*j*_ is a random variable (e.g. through dependence on time, temperature, or other measured quantities treated as random variables), then the variance of the probability of pathogen spillover, *P*_*S*_, can be decomposed into the sum of variances and covariances of log-probabilities, log(*p*_*j*_),

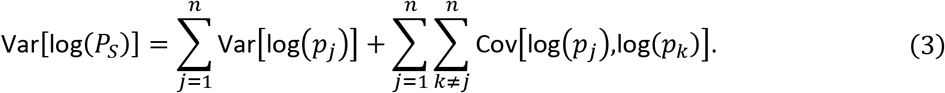

If any two events have a perfect, negative correlation on a log scale, they won’t contribute to variance in the overall probability of spillover. All else being equal, if changing the ambient temperature halves the probability a virus survives long enough to contact a but doubles the probability of infection given contact, the risk of pathogen spillover will remain unchanged human [9, 10, 11, 12]. More generally, the covariances between events determine whether a measured change in any given log(*p*_*j*_) will have an equal, dampened, or amplified effect on the overall probability of spillover. Pathogen spillover is an inextricably multilevel process: one cannot make inferences about how predictable changes in the probability of success at one level, such as how temperature is associated with the duration of environmental persistence, affect the overall probability of pathogen spillover without knowing concomitant changes in the probability of success at every level, such as how temperature is associated with shedding, contact, and the probability of infection. The only way a measured change in a log(*p*_*j*_) produces an equal change in log(*P*_*S*_) is if one assumes or knows that all other levels remain unchanged.

The log decomposition of variance in the probability of pathogen spillover produces some useful concepts for discussing the multilevel modeling and management of pathogen spillover. First, there may be an unknowable total variance in spillover risk as we can’t measure Var[log(*P*_*S*_)]. As studies accumulate statistical associations between environmental covariates and log probabilities of success, log(*p*_*j*_), some variance in pathogen spillover risk is observed and explained. Not all variance explained by environmental covariates may be accessible through management interventions, and so an important terminological distinction is that between “explained variance” and “manageable variance” (Box 1). Manageable variance is the overall variance in spillover risk which can be modulated through allowable management interventions or a set of management interventions under consideration [Sokolow et al., this issue].

Analogous concepts can be derived using the derivative of the vector of log-probabilities, 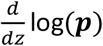, to parameterize the directional change in log probabilities of spillover resulting from changing environmental covariates or management actions defined by a variable *z*. For example, if *z* is temperature, then we can model temperature-dependent probabilities of success as ***p***(*z*). Noting that

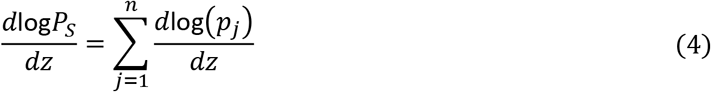

one obtains the net effect of the management action or environmental covariate on spillover risk by projecting the vector 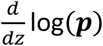 onto the one vector, **1**. If the inner product 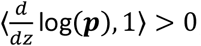, then the risk of pathogen spillover will increase with increasing *z*, and isoclines of *P*_*S*_ are linear subspaces: planes of log(***p***) orthogonal to 1. The linearity log(*P*_*S*_) as a function of each level’s log probability is mathematically useful to note (insofar as the underlying model pertains to reality): linearity permits simple dimensionality reductions by decomposing changes in log(***p***) into changes along the simplex-like isoclines on which spillover risk is constant and changes along the single direction along which spillover risk increases or decreases [13].

Analysis of a serial percolation process for pathogen spillover yields an intuitive scaffold for managers attempting to make decisions based on literature from multiple levels: the net impact of a covariate or management action depends on an unavoidably multilevel process. Only when the impact of a management lever [Sokolow et al., this issue] on probabilities of pathogen success across many levels is known (or justifiably assumed) can one estimate the impact of management actions on pathogen spillover risk. Such estimations can be approximated by considering log-scale changes in the probability of various events on the pathway to spillover occurring.

### Alternative pathways / Detours

In many cases, the assumption of a single pathway to pathogen spillover may be too restrictive. For example, a pathogen like Bartonella could be transmitted by one of many transmission routes [14, 15]. In these scenarios, the graphical model becomes

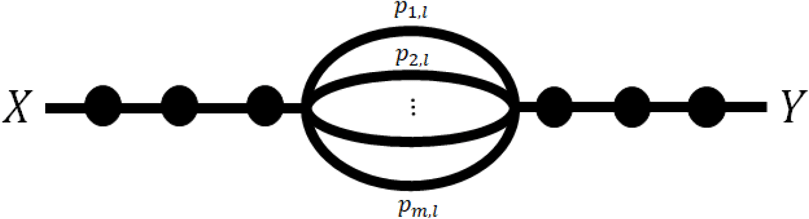

where level *l* has *m* alternative pathways (in the case of Bartonella, alternative transmission routes) and *p*_*i,l*_ is the probability of a pathogen surviving the *i^th^* alternative pathway given entry to that pathway. Letting *α*_*i*_ be the fraction of particles in the pool prior to *l* which go down the *i^th^* alternative pathway such that Σ_*i*_*α*_*i*_ = 1, the probability a given particle spills over becomes

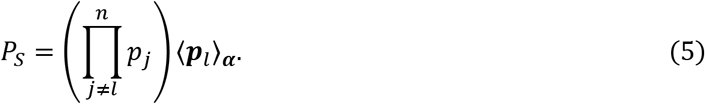

where ⟨***p**_l_*⟩_***α***_ = Σ_*i*_ *α*_*i*_*p*_*i,l*_ is the weighted average probability of surviving level *l*.

Taking the logarithm of both sides of equation (5), we encounter a nonlinearity through the logarithm of an arithmetic mean,

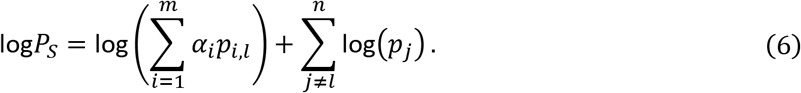

This log-sum nonlinearity affects our ability to make neat inferences on the impact of doubling or halving the probabilities of success along one of the alternative pathways. Similarly, the variance of a log-sum cannot expressed in terms of variances and covariances of each log-probability. Taking the derivative of equation (6) with respect to a covariate, *z*, yields

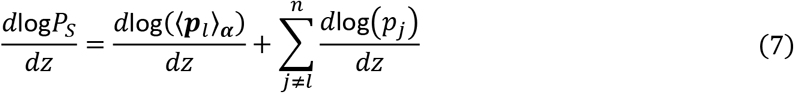

and, noting that,

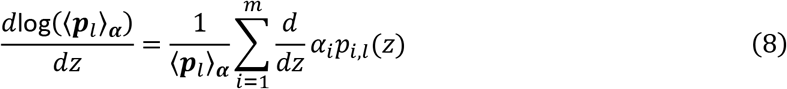

we see the impact of unit changes on one alternative pathway can’t be separated from the probabilities of other alternative pathways due to the inverse dependence on the mean, ⟨***p**_l_*⟩_***α***_. The analysis becomes even more complicated if the fraction of pathogens going down alternative pathways, ***α***(*z*), depends on the covariate *z*. Within alternative pathways, equal and opposite arithmetic changes in edge percolation probabilities, *α*_*i*_*p*_*i,l*_, as opposed to log-scale changes, will cancel out.

While there are analytical tools for variance decomposition through Taylor Series approximations of the log-sum function, that is beyond the scope of this paper. Instead, we want to emphasize that the simplest deviation from the serial percolation model produces a log-sum function in equation (6) that prevents the immediate generalization of the variance decomposition from the serial model. The log-sum will come up again. Since the operands of the log-sum are all positive, they can be represented as an exponent, 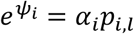, yielding the LogSumExp function, also known as the softmax function because

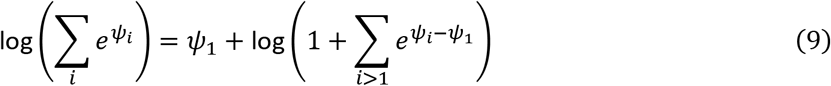

is approximately equal to the maximum, *ψ*_1_ = max{*ψ*_*i*_}, when *ψ*_1_ is sufficiently prominent (i.e. 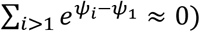. If *ψ*_1_ is the alternative pathway with the most percolation of pathogens, what we refer to as the “dominant pathway”, then the percolation through all of the alternative pathways, including its associations with covariates, will resemble percoluation through the dominant pathway in a manner given by the softmax function. We further discuss and demonstrate the role of the softmax function on parameter estimation in the section on statistical inference.

### Alternative Sources

Many pathogens can spill over from one of multiple reservoirs. Examples of pathogens spilling over from multiple animal reservoirs include plague [16], *E. coli* [17], giardia [18], Crypotospordiosis [18], Lyme disease [19] and more. For *m* alternative reservoirs whose shed or vector-borne pathogens follow independent pathways until a common pool at level *l*, the percolation graph becomes

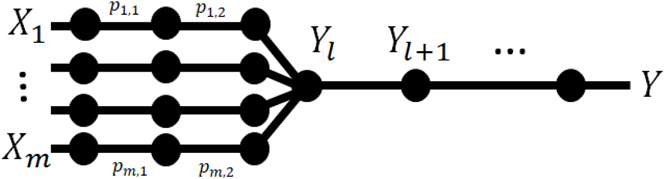

where *X_i_* is the amount of pathogen released from reservoir *i, p*_*i,j*_ is the probability of the pathogen released from reservoir *i* successfully passing level *j*, and *Y_l_* is the amount of pathogen pooled at step *l*, after *l* logical events. We will use *l* to denote the first point at which pathogens from different reservoirs are in a common pool, such as influenza virions shed into a pond by waterfowl [20, 21].

Prior to pooling, each pathway is a separate, serial percolation process with the usual results: a Poisson random variable at the shedding process for species *i, X_i_*~*Pois*(*λ_i_*) yields a filtered Poisson random variable prior to pooling denoted *Y*_*i,l*–1_~*Pois*(*λ_i_* Π_*j*<*l*_ *p*_*i,j*_). *Y*_*i,l*–1_ is a filtered Poisson because it can be expressed as a Poisson random variable for which counts are subsequently passed through a Bernoulli filter with success probability Π_*j*<*l*_ *p*_*i,j*_; virions released from a Poisson shedding process be removed with probability 1 – Π_*j*<*l*_ *p*_*i,j*_.

At the intersecting node of the pathways from alternative sources, a set of Poisson random variables are added to yield the number of pathogens in the common pool having survived a series of *l* events. Since the sum of Poisson random variables is a Poisson random variable with rate equal to the sum of the rates, the number of pathogens surviving up to the common pool will be

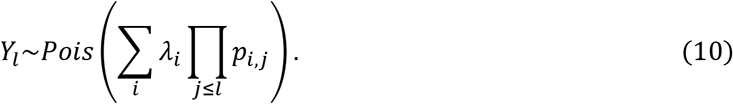

The number of spillover events can be calculated by recognizing that percolation is serial between *Y_l_* and *Y* (table 1). For pathogens spilling over from multiple sources each shedding a Poisson-distributed number of pathogens, the pooled number of spillover events from all sources will be a Poisson random variable with a rate equal to the sum of other rates. Such pooling is commonly modeled and analyzed in queuing processes, a conceptual connection we save for future work.

### Time-Dependence – percolation processes

In all the discussed different probabilistic model structures for pathogen spillover, we’ve implicitly assumed an instantaneous count process – Poisson or negative binomial – generating a pulse of pathogens and an instantaneous filtration of the shed pathogens to yield similar count processes at each point on the pathway to spillover. Often, however, shedding is a time-dependent process, and our data (e.g., the number of bird flu virions found in a pond) consist of a sampling of pathogens that have survived up to a step in the spillover pathway at a point in time. In addition to shedding, the survival probabilities of a pathogen may depend on time, such as seasonal patterns in temperature affecting the rate of decay of pathogens in the environment.

Let *λ*(*t*) be the propensity of viral shedding at time *t*. Consider the time-dependence of environmental persistence combined with a variable time, *τ*, between shedding and contact. For pathogen spillover to occur, viruses shed into the environment must survive until contacting a human. The environmental pool at time *t, Y*_*l*_(*t*), will contain viruses shed at various times in the past that have survived up to time *t*. Let *τ~h*(*τ*) be a random variable with density *h, p*_*l*_(*t, τ*) be the probability of a pathogen surviving in the environmental pool from time *t* − *τ* to *t*, and assume that all previous logical events partitioned are instantaneous but with time-dependent probabilities, *p*_*j*_(*t*). Then, *Y*_*l*_(*t*)~Pois(*λ*_*l*_(*t*)), where

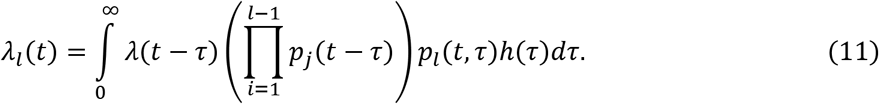

The rate defined in equation (11) is equivalent to the model of alternative sources in equation (10) with appropriately shifted time-dependent probabilities. Hence, nonstationary probabilities of passage and random time intervals for lagged passage through a particular level produce a percolation model of alternative sources in time for all steps including and prior to the lagged step, as illustrated below, where darker shading is used to indicate more recent shedding times.

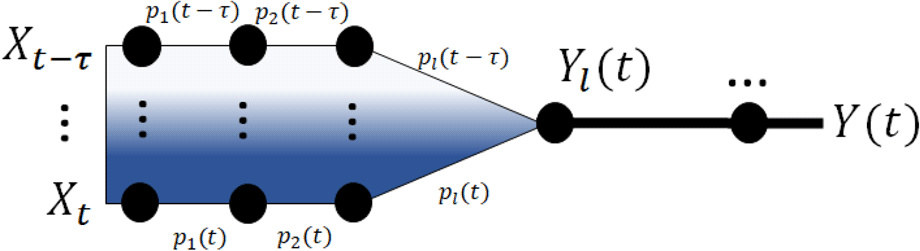

Stochastic epizootiological and shedding processes can be integrated over time to produce models of spillover risk by employing a formally defined birth-death process with inhomogeneous hazard rates *λ*(*t*) [22] as approximated with a Gillespie algorithm [23]. Keeping track of the hazard rate allows one to use the analytical results from alternative sources, in particular the softmax nonlinearity, to understand how inferred variation in the overall spillover risk is driven by variation in the maximum-risk pathways in time.

## Statistical Inference

With the inextricably multilevel nature of pathogen spillover, one aim is to estimate the relative importance of various levels in the pathway to spillover using data on the incidence of pathogen spillover [24, 25, 26]. Barring data on intermediate pools of pathogens *Y_j_*, one may hope to measure relative importance of each level through the curation of a dataset with level-specific covariates, *z_j_*, hypothesized to explain some variation in the probability of spillover for level *j*. For example, in an alternative source model one may measure the population densities of all alternative sources in the locales where spillover occurred. Typically, large-scale studies of spillover work with a set of pooled counts, *Y*, representing successful infections at the end of the percolation model of spillover, and we hope to use regression – often generalized linear or additive modeling – to determine spillover risk and assess the relative importance of different covariates and barriers to spillover.

Percolation models easily demonstrate two challenges for such statistical inferences using pooled counts such as the number of spillover events. The challenges arise from the difference between the probability and rate parameters used in the derivation of equations 1–11, and the canonical parameters, logit probabilities and log-rates, used for generalized linear modelling of exponential family random variables. While the probability and rate parameters combine nicely for tractable analyses of percolation models, statistical inference through GLMs frequently uses canonical parameters obtained by evaluating nonlinear link functions of the probability or rate parameters. The nonlinearity of link functions for the Poisson, negative binomial, and Bernoulli random variables present in the percolation models of spillover have important consequences for interpreting results from regression on *Y* and the inference of the relative importance of various steps and pathways to spillover. In this section, we will denote canonical parameters *η*(*z*) as functions of a single covariate *z*, and the random variable for that covariate will be *Z*.

### Serial

The simplest percolation model of pathogen spillover for statistical inference will contain a pathogen pulse, *X*~*Pois*(*λ*(*z*)), and the probability of spillover, *P_*s*_*(*z*), combined to yield the number of spillover events, *Y*~*Pois*(*P_*S*_*(*z*)*λ*(*z*)). In a GLM framework, one would model the canonical parameters of the two processes – counts of virions shed from reservoirs and the probabilities of survival through a percolation process – separately as

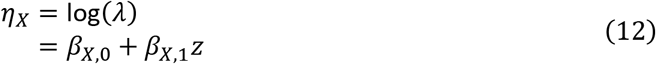

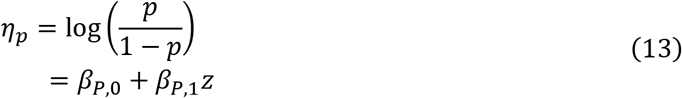

However, a GLM for spillover events, *Y*, would estimate *η*_*Y*_, the log of the rate for *Y*. Using the rate of *Y* defined by the percolation process, combined with equations (12) and (13), we see that a GLM predicting *Y* would model

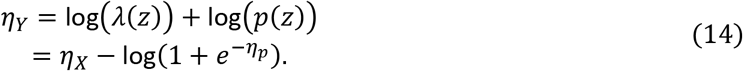

Substitution with equations 12 and 13, yields

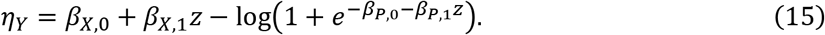

If *η*_*p*_ ≪ 0, the approximation 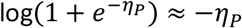 permits approximate superposition of intercepts and slopes for an overall linear model, yielding

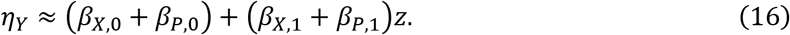

However, if *η*_*p*_ ≫ 0, the approximation 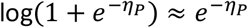 produces terms which are nonlinear in *z*_*j*_, yielding

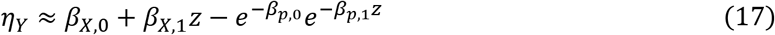

These approximations and the switching points between them near *η*_*Ρ*_ = 0 are illustrated in Figure 2.

**Figure 2:**
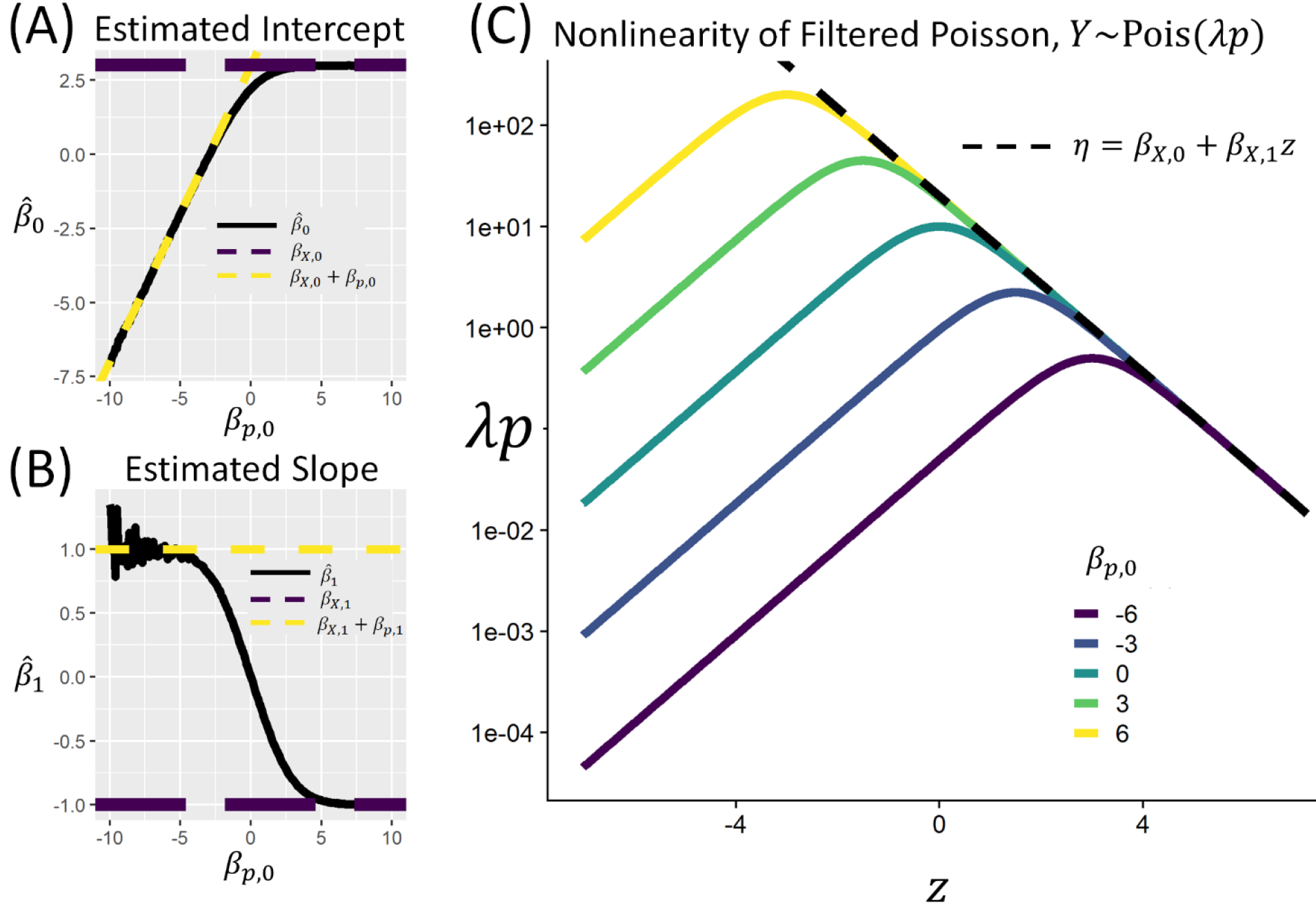
Simulations confirm the nonlinearities of equations (13) and (15). (A-B) The nonlinearity is well approximated with a linear model when attrition rates are high, *η*_*Ρ*_ ≫ 0, or when attrition rates are low *η*_*p*_ ≪ 0. By simulating 1 × 10^5^ Poisson random variables, varying *β*_*p*,0_ while fixing *β*_*Χ*,0_ = 3, β_*X*,1_ = –1 and *β*_*p*,1_ = 2, we see the convergence of estimated intercepts and slopes to that predicted by equations (13) and (15). (C) Using the same parameterization and plotting *λ*_*p*_ as a function of *z* on a log scale illustrates the switching behavior of the nonlinearity in *η* = log(*λp*). For sufficiently large *z, η*_*p*_ ≫ 0 and the approximation *η* ≈ *β*_*Χ*,0_ + *β*_*Χ*,1_*z* holds (dashed line). The potential for such nonlinearities in real world data motivate the use of nonlinear models, such as GLMs or the explicit nonlinear model defined in equation (19).

The nonlinearity of canonical parameters as functions of rate and probability parameters in percolation processes yields one important caveat of statistical inference of the association between covariates and the number of spillover events. Where the datasets and covariates are limited to observing high probabilities of pathogen success given shedding (*P_S_* ≫ 0.5), estimates of regression coefficients will resemble the associations driving shedding (*η*_*X*_), causing one to underestimate the potential importance of attrition when projecting beyond observed data to extreme but feasible covariate values yielding *p* ≪ 0.5. Equation 14 shows how percolation models produce tractable and sometimes linear combinations of the parameters describing the percolation process’s input rates and probabilities, yet nonlinear functions of canonical parameters typically used in GLMs of a single level. Consequently, one may assume a GLM for each level or a GLM for the number of spillover events, but not both.

Consider, for example, a carefully curated dataset of level-specific data, *z_j_*, such that

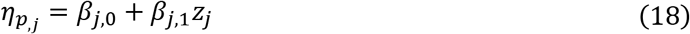

for levels *j* = 1,…, *n*, and shedding having a canonical parameter dependent upon *z_X_* as modeled as in equation 12. The number of spillover events will then have canonical parameter

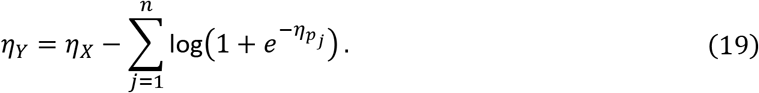

If over the observed values of {*z*_*j*_}, levels can be summarized as either strong attritions with *η*_*p,j*_ ≪ 0 for all *j* ∈ *S* or weak attritions *η*_*p,j*_ ≫ 0 for all *j* in *W*, where *S* ⋃ *W* = {1,…, *n*} captures all the logical events between shedding and spillover, then approximations used for equations 16 and 17 yields

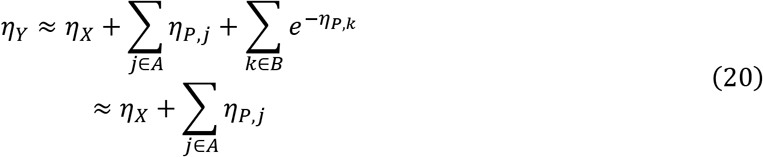

Levels with weak attrition rates in the observed data – even if they are more sensitive to feasible changes in covariates – will be nonlinear, and their importance as manageable levels will be underestimated as they contribute little to the canonical parameter for the overall rate of spillover. If one agrees with a percolation model for spillover, then GLMs should not be used to estimate relative importance of various levels. Unless estimating the correct nonlinear model in 18, one should be more wary than usual of projecting risk for covariate values far beyond those used to train a model (Figure 2).

### Alternative Pathways and Sources

As we show above, the number of spillover events for alternative pathways and sources, whether alternative reservoirs or alternative sources in time, will be a Poisson random variable whose canonical parameter will include a softmax function of the rates or probabilities for alternate pathways to spillover (equation 9). The softmax nonlinearity provides some insight into how investigations based on GLMs of spillover risk can err. If *Y*_*i*_~Pois(*λ*_*i*_) for alternative pathways 1,…, *m*, then the pooled number of spillover events, *Y* = Σ_*i*_*Y*_*i*_, is a Poisson random variable *Y*~Pois(Σ_*i*_*λ*_*i*_). The canonical parameter one would estimate for GLMs of *Y* will be

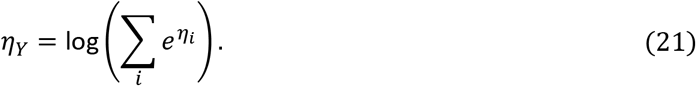

The softmax functional form of *η*_*Y*_ provides some insight about its behavior. First, *η*_*Y*_ will behave more like the maximum of {*η*_*i*_} as the proportion of spillover events arising from the dominant pathway increases. Consequently, GLMs of pooled counts or spillover events across alternative sources will produce regression coefficients reflecting those of the maximum-risk i.e. dominant pathway (Figure 3).

**Figure 3.**
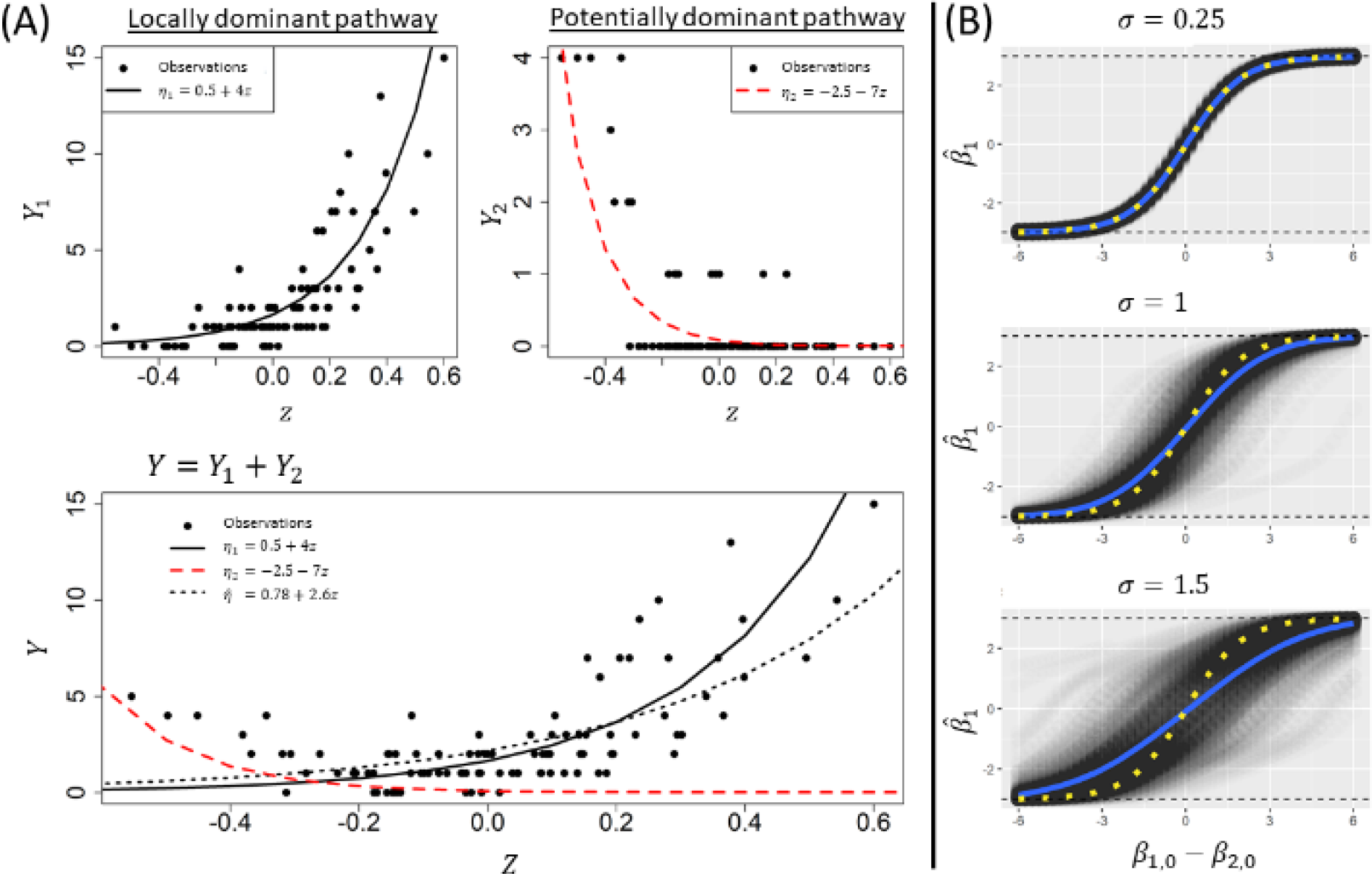
Alternative pathways, whether across reservoirs or across time, result in a nonlinearity in the canonical parameters. If *Y*_*i,l*–1_~Pois(*λ*_*i*_) is the pathogen contribution from alternative sources *i* = 1, …, *m*, then the pool of pathogens contributing to spillover, *Y*_*l*_~Pois(Σ_*i*_*λ*_*i*_) has canonical parameter 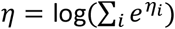 where *η*_*i*_ = log(*λ*_*i*_). This nonlinearity, the LogSumExp or softmax function, biases linear regression coefficients, including slopes, towards the pathway with the highest mean or intercept, what we refer to as the dominant pathway. (A) The bias of the softmax function for linear regression is shown. If two pathogens have different regression coefficients with a standard Gaussian covariate, *z*~Gsn(0, *σ*^2^), but one has a larger mean under the observed covariates, the pathogen with the larger mean will dominate linear regression. Consequently, generalized linear models of pathogen spillover will underestimate the risk of spillover when an alternative pathway becomes dominant. Such effects can reduce the effectiveness of multiple regression efforts to predict spillover risk. Generalized additive models or regression clustering may alleviate this issue (B) Linear regression of pooled Poisson random variables with different covariate regression coefficients, *β*_*i*,1_, will exhibit threshold switching towards the regression coefficient of the dominant pathway. The estimated slope is reasonably approximated by a weighted average of the slopes, with weights equal to the mixing proportions of pathogens in the pool (yellow dots).

Second, if linear models for each pathway are the correct model for each level but the maximum-risk pathway is globally dominant within a dataset (e.g., if different reservoirs are dominant in different geographic regions or different time periods), a linear model of *Y* will fit a plane, 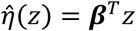, across a convex bowl

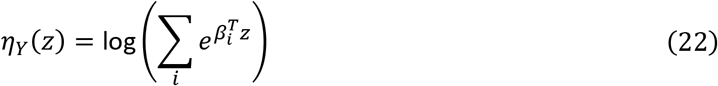

leading to a potentially interpretable nonlinearity (Figure 3a). While one should always look for heteroskedasticity to ensure a model is a good fit, it’s especially important to check when predicting pathogen spillover risk from alternative sources in space, time, or species, as overdispersion could suggest either nonlinearity in the constituent canonical parameters or linearity in the constituent canonical parameters for each level combined with switching between dominant pathways of spillover within a dataset. Heteroskedasticity does not imply reservoir switching, but spillover data analysts should be extra wary because pathway switching is an additional potential source of heteroskedasticity in non-serial percolation models. Regression clustering or switching-regression models [27, 28] may separate data points with different dominant pathways for overdispersion due to pathway switching; this could be a fruitful avenue for future research.

## Conclusions

Pathogen spillover is an inherently random, multilevel process [3]. Here, we have developed the inherently random and multilevel percolation-based models of pathogen spillover. Percolation models can be visualized with graphs indicating our mental model of the pathways to spillover, and various percolation models can be analyzed to yield clear connections between model structure, data analysis, and management paradigms for inferring and mitigating spillover risk.

For a serial percolation model, the log-probability of pathogen spillover allows one to decompose variance in spillover risk in terms of variances and covariances of log-probabilities of each level, providing a basic multilevel model of pathogen spillover. The serial decomposition of spillover risk may be of use for those developing statistical and mathematical methods to model and infer spillover risk. The serial model of pathogen spillover produces some conceptual tools, illustrated in Figure 1, towards a management paradigm to evaluate the relative costs and benefits of various management actions, the information gain of data from different levels through increased explained variance in log-probabilities of success, and more.

We have shown how percolation-based models lend themselves to easy visualizations of model structure and tractable calculations of spillover rates. In addition, we have provided two important results for the inference of spillover risk across levels and across alternative pathways. In general, one cannot use GLMs for both levels or alternative sources and the overall rate of spillover. While probabilistic percolation-based models of the number of spillover events yields count distributions which are easily described as products of probabilities and sums of rates, such quantities are nonlinear functions of the canonical parameters used for the individual levels.

Consequently, despite the demand for inferences of relative importance of various pathways in spillover and the ease of inferring relative importance in generalized linear models, we strongly discourage the default use of GLMs predicting the number of spillover events. For alternative source models, we hypothesize that regression clustering algorithms may allow one to separate out the linear models of alternative levels. When using a GLM to predict pathogen spillover risk, it’s especially important to check for heteroskedasticity as bowl-shaped heteroskedasticity may indicate a meaningful and consequential alternation of dominant pathways to spillover within the dataset. Instead, generalized additive models or the appropriate nonlinear models specified in equations 20 and 22 should be estimated to appropriately estimate attrition rates at various levels in the pathway to spillover.

Percolation models will fail when virion replication in intermediate pools is non-negligible, such as replication within alternative pathways reliably leading to one virion causing potentially multiple infections. Similarly, percolation models do not capture epizootiological feedbacks which may exist between environmental or vector pools and wildlife reservoirs. Finally, where dose-response filters are crucial, the spatial distribution of virions may be relevant and will require additional model complexity.

Percolation-based models capture the minimal assumptions of pathogen spillover and allow a tunable level of complexity for modeling spillover risk ranging from a coin toss to a fully deterministic system in which alternative sources and pathways are determined with probability 1. Percolation pipelines can also be a modular component for models of spillover risk. One could model shedding as an externally defined stochastic process, as we have assumed here, or one could model shedding into a percolation pipeline through more commonly employed SIR-type epizootiological models. Even deterministic models can be input into percolation pipelines, as a posterior distribution over different model structures can be incorporated through a stochastic sampling of different m°de\s’ shedding trajectories over time or space. All models are wrong, but percolation models may be a useful tool for conceptualizing, managing, and analyzing the risk of pathogen spillover.

## Supplemental calculations

### Proof of Poisson & Negative Binomial stability to binomial filtration

***If X~Pois*(*λ*) *and Y~Binom*(*X,p*), *then Y~Pois*(*λp*).**

By the law of total probability, the probability mass function of *Y* is

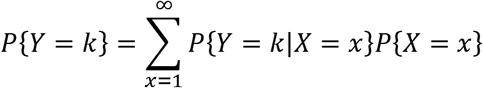

Noting that *P*{*Y* = *k*|*X* = *x*} = 0 for *x* < *k* and substituting the binomial and Poisson probability mass functions, we get

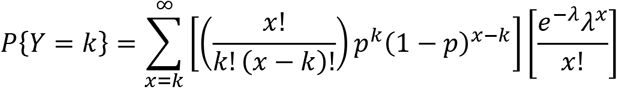

Substituting *m* = *x* – *k* and combining a few terms, we get

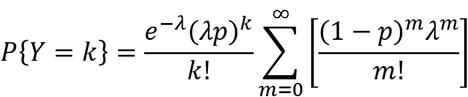

Noting that the infinite sum is the Taylor expansion for *e*^*λ*(1–*p*)^, we get

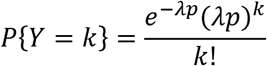

QED.

***If X~NegBinom*(*r, q*) *and Y~Binom*(*X, p*), *then* 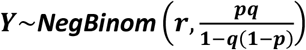.**

Repeating the first two steps above, but for a negative binomial probability mass function for *X*

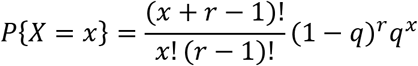

yields

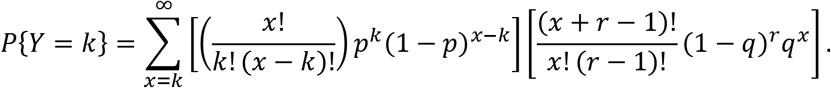

Cancelling and collecting a few terms, and using the same substitution *m* = *x* – *k*, we get

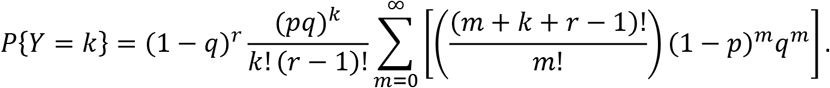

Multiplying and dividing by (*k* + *r* – 1)! Yields

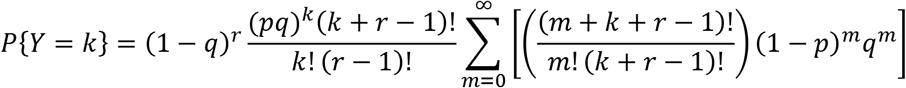

Which can be written more simply using the binomial coefficient

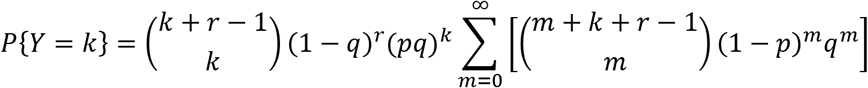

Multiplying and dividing by (1 – *q*(1 – *p*))^*k*+*r*^, and moving the term (1 – *q*(1 – *p*))^*k*+*r*^ inside the summand yields

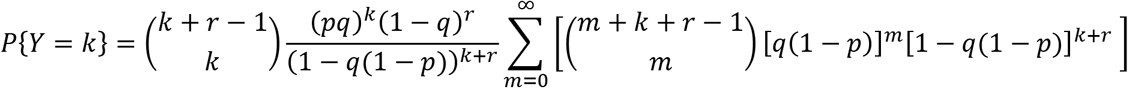

The summand is the probability mass function of a negative binomial random variable with *k* + *r* failures and success probability *q*(1 – *p*). As such, the infinite sum is equal to 1 and we’re left with

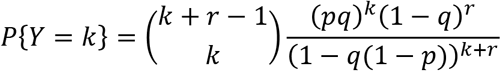

Combining terms yields the familiar probability mass function

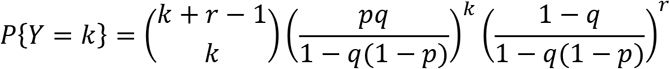

For a negative binomial, therefore

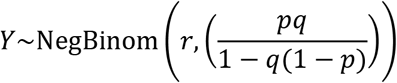

QED.

***Parameterized by mean μ and dispersion ψ: If X~NegBinom*(*μ,ψ*) *and Y~Binom*(*X,p*)*, then Y~NegBinom*(*μp,ψ*).**

Using the formulas for the mean 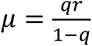 and dispersion 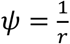, we note that the dispersion of *X* and *Y* are equal, while the mean of *Y*, denoted *μ*_*Y*_, becomes

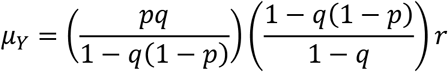

which simplifies to

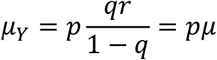

QED.

